# *Keetia tinka* sp. nov. & *K. kounounkan* sp.nov. (Rubiaceae - Vanguerieae) new threatened forest climbers and shrubs of sandstone plateau of the Republic of Guinea

**DOI:** 10.1101/2025.08.27.672669

**Authors:** Faya Julien Simbiano, Charlotte Couch, Sekou Magassouba, Jack Plummer, Martin Cheek

## Abstract

Two new species of *Keetia* are described from recent botanical collections for conservation management made in surviving submontane forest areas of the sandstone plateau areas of the Republic of Guinea. *Keetia kounounkan* Cheek & Simbiano is a shrub of the Kounounkan Plateau towards the border of Sierra Leone, so far with a single location in gallery forest. *K. tinka* Cheek & Simbiano is an evergreen rainforest climber of the main part of the Fouta Djalon Highlands with two locations both in degraded forest.

Both species are described, illustrated and provisionally assessed for their conservation status, the first as Critically Endangered, the second as Endangered.

## INTRODUCTION

*Keetia* E. Phillips (Rubiaceae, Vanguerieae) was segregated from *Canthium* Lam. by Bridson (1985, 1986). This genus of about 41 accepted species (Cheek & Onana 2024) is restricted to sub-Saharan Africa (excluding Madagascar and the Mascarene Islands) and extends from Senegal and Guinea in West Africa (Gosline *et al*. 2023a; 2023b) to Sudan (Darbyshire *et al*. 2015) in the North and East, and S. Africa in the South (Bridson 1986). *Keetia* differs from other African genera of Vanguerieae by its pyrenes with a fully or partly-defined lid-like area around a central crest, and endosperm with tanniniferous areas (Bridson 1986). *Keetia* species are usually climbers (very rarely shrubby) and occur mostly in forest habitats, but sometimes in wooded grassland. In a phylogenetic analysis of the tribe based on morphology, nuclear ribosomal ITS and chloroplast *trnT-F* sequences, Lantz & Bremer (2004), found that based on a sample of four species, *Keetia* was monophyletic and sister to *Afrocanthium* (Bridson) Lantz & B. Bremer with strong support. Highest species diversity of *Keetia* is found in Cameroon and Tanzania, both of which have about 15 taxa (Onana 2011; POWO, continuously updated) and in Gabon, where 10 species are currently recorded (Sosef *et al*. 2006) but around 25 are actually present, many of them undescribed (Lachenaud pers. comm. to Cheek, 2024). Recently, bacterial leaf nodulation was discovered to occur in the genus, only the fourth genus of the family in which this is recorded (Cheek & Onana 2024). Several *Keetia* species are point endemics, and have been prioritized for conservation (e.g. Onana & Cheek 2011; Couch *et al*. 2019; Murphy *et al*. 2023) and one Guinean threatened species, *Keetia susu* Cheek has a dedicated conservation action plan (Couch *et al*. 2022).

Bridson’s (1986) account of *Keetia* was preparatory to treatments of the Vanguerieae for the Flora of Tropical East Africa (Bridson & Verdcourt 1991) and Flora Zambesiaca (Bridson 1998). Pressed to deliver these, she stated that she could not dedicate sufficient time to a comprehensive revision of the species of *Keetia* outside these areas: “full revision of *Keetia* for the whole of Africa was not possible because the large number of taxa involved in West Africa, the Congo basin and Angola and the complex nature of some species would have caused an unacceptable delay in completion of some of the above Floras. […] A large number of new species remain to be described.” (Bridson 1986). Several of these new species were indicated by Bridson (1986), and other new species by her arrangement of specimens in folders that she annotated in the Kew Herbarium. One of these species was later taken up and published by Jongkind (2002) as *Keetia bridsoniae* Jongkind. In the same paper, Jongkind discovered and published *Keetia obovata* Jongkind. Based mainly on new material, additional new species of *Keetia* have been published by Bridson & Robbrecht (1993), Bridson (1994), Cheek (2006), Lachenaud *et al*. (2017), Cheek *et al*. (2018a), Cheek & Bridson (2019), Cheek & Onana (2024), Cheek et al. (2024a), Cheek *et al*. (2024b) and there are several other specimens that fit no other species, (e.g. Cheek *et al*. 2004; 2011) and remain to be described.

In this paper we continue the project towards an updated taxonomic account of *Keetia* by describing from recently collected material two further new species from Guinea, *K. tinka* Cheek & Simbiano (previously considered to be a variant of *K. magassoubiana* Cheek) and *K. kounounkan* Cheek & Simbiano (initially identified as *K. susu*).

In recent years, numerous new species to science have been described from Guinea, such as from the surviving remnants of species-diverse forests. These include species of climbers e.g. in *Monanthotaxis* Baill. (Annonaceae, Hoekstra *et al*. 2021), *Hibiscus* L. (Malvaceae, Cheek *et al*. 2020a), *Keita* Cheek (Olacaceae, Cheek *et al*. 2024c), small trees and shrubs e.g. *Casearia* Jacq. (Salicaceae, Breteler & Baldé 2024), *Tarenna* Gaertn. (Rubiaceae, Jongkind 2021), and *Tabernaemontana* L. (Apocynaceae, Jongkind & Lachenaud 2022), non-chlorophyllous heteromycotrophs (*Gymnosiphon* Blume, Burmanniaceae, Cheek et al. 2024d) and from waterfalls, rheophytes e.g. *Inversodicraea* Engl. and *Saxicolella* Engl. (Cheek *et al*. 2017; 2022). These discoveries are set to continue so long as funds to support taxonomists and taxonomic work continues and habitat survives.

## MATERIALS AND METHODS

Names of species and authors follow IPNI (continuously updated) and nomenclature follows Turland *et al*. (2018). Herbarium material was collected using the patrol method e.g. Cheek & Cable (1997) and processed and studied as in Davies et al. (2023). Herbarium specimens were examined with a Leica Wild M8 dissecting binocular microscope fitted with an eyepiece graticule measuring in units of 0.025 mm at maximum magnification. The drawing was made with the same equipment with a Leica 308700 camera lucida attachment. Pyrenes were characterized by simmering selected ripe fruits in water until the flesh softened and could be removed by scalpel. A toothbrush was then used to clean the pyrene surface to expose the surface sculpture and the lid. Finally, a fine saw was used to cut a transverse section of the fruit and seed, allowing observation of tanniferous cells in the endosperm and measurement of the endocarp thickness. Specimens were inspected from the following herbaria: BM, FHO, HNG, K, P, SL and YA and images of specimens on Gbif.org.

Google Earth Pro was used to view the collecting sites, read accurate elevations, to assess the continued survival of the species using as proxy the continued existence of forest habitat at the collection site, and also to evaluate likely extent of occurrence sensu IUCN (2012) for the conservation assessment. The format of the description follows those in other papers describing new species of *Keetia*, e.g. Cheek *et al*. (2025). Terminology follows Beentje & Cheek (2003). All specimens indicated “!” have been seen. The conservation assessment follows the IUCN (2012) standard. Herbarium codes follow Index Herbariorum (Thiers, continuously updated).

### TAXONOMIC TREATMENT

The first of the two new species, *Keetia kounoukan*, had been initially identified as the locally more frequent *K. susu* to which it is superficially similar, also being a large-fruited shrub or small tree of sandstone habitats in Guinea. However, on closer examination it was found to differ in so many unusual character traits that it is not clear with which species in the genus its closest affinities with (Table 1 below). In the key to the *Keetia* species of West Africa (Cheek et al. 2025) it fits neither of the leads in the first couplet, having patent brown hairs on the stem > 0.5 mm long. Although the fruits resemble those of *K. susu* and its allies, they lack the greatly accrescent disc, and the pyrene lid and sculpture are completely different (Table 1).

**Table 1.** Selected diagnostic characters separating *Keetia kounoukan* from *K. susu*.

|  | *Keetia susu* | *Keetia kounoukan* |
| --- | --- | --- |
| Petiole length and indumentum | 4–16 mm, glabrous | 5–10 mm, densely hairy |
| Leaf acumen apex | Acute | Rounded |
| Domatia | Pit domatia (hairs inside pit) | Tuft domatia (hairs exserted) |
| Fruit size and colour | 13–17 × 15–20 × 11–13 mm, glossy black when ripe | 10 × 11–12 × 6–7 mm, matt brown when ripe |
| Fruit disc | 5–8 mm in diameter | 1–3 mm in diameter |
| Pyrene lid and pyrene surface sculpture | Lid ventral, lacking crest; pyrene surface with irregular raised areas. | Lid apical indistinct, crest with midline furrow; surface smooth, with fingerprint pattern (Fig 1K) |
| Stipule awn length | 4–4.5 mm | 4–8 mm |
| Stem habit, indumentum | Primary stems scandent, some secondary shoot pairs reflexed; glabrous | Primary stems not scandent, secondary shoots ascending, densely hairy (distal internodes). |

***Keetia kounounkan*** *Cheek & Simbiano* ***sp. nov***. Type: Guinea, Forécariah Prefecture, southern plateau of Kounounkan Massif, 9° 33’ 01.6” N 12° 50’ 20.4” W, 1100 m, fr., 5 Feb. 2019, *van der Burgt 2262* with P.M. Haba, Konomou & Xanthos (holotype K! barcode K001971152; isotypes BR, HNG barcode 0002731). Fig. 1-3

**Fig. 1.**
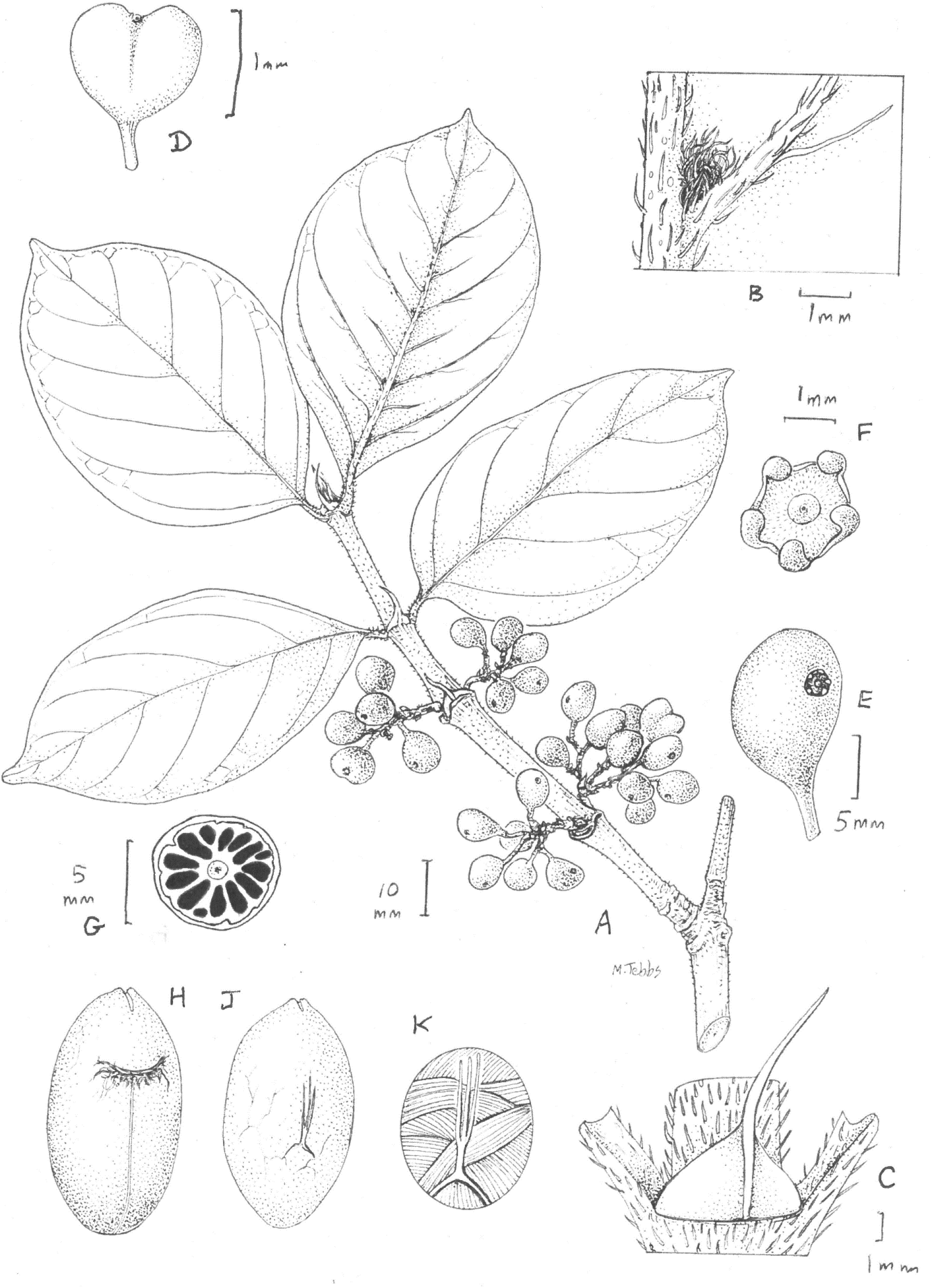
Keetia kounounkan. **A** secondary stem with infructescences; **B** domatium, on lower leaf surface; **C** stipule; **D** 2-seeded fruit; **E** 1-seeded fruit; **F** disc and calyx of a 1-seeded seed; **G** transverse section of seed showing tanniniferous rays; **H** pyrene ventral view; **J** pyrene dorsal view; **K** detail of surface of pyrene. All from *van der Burgt* 2262 (K). Drawn by Margaret Tebbs.

#### Diagnosis

*Keetia kounounkan* is similar to and was initially identified as *Keetia susu* Cheek but *K. susu* has a glabrous petiole, (the petiole of *Keetia kounounkan* is densely hairy). The leaf acumen of *K. susu* is acuminate, with a sharp point, while in *K. kounounkan* the acumen has a rounded apex. The domatia of *K. susu* are pit domatia, hairy within the pit only, while the domatia of *K. kounounkan* are tuft domatia with exserted hairs. The fruits of *K. susu* are larger, 13–17 × 15–20 × 11–13 mm, glossy black when ripe, while the fruits of *K. kounounkan* are smaller, 10 × 11–12 × 6–7 mm, brown when ripe. The fruit disc of *K. susu* is 5–8 mm in diameter, while the fruit disc of *K. kounounkan* is 1–3 mm in diameter. See Table 1 for additional diagnostic characters.

#### Description

Shrub, 2 m high, 4 m wide, stem to 5 cm thick at base. Primary stems erect, not climbing, secondary shoots ascending, stout, bearing usually two pairs of leaves at stem apex, fruiting from leafless nodes. Leafy stems cylindrical, drying grey to black, internodes 2–4.5 cm long, lenticels inconspicuous, young stems densely hairy, hairs simple, persisting to the third node from the apex, brown, straight, stout, acute, appressed to subappressed, 0.2–0.9 mm long, extending to the petiole, abaxial midrib, secondary veins, margins of leaf-blades, and the infructescence axes, older stems glabrescent. Stipules persistent to the third node, glabrescent, 7–12 mm long, base broadly triangular, 2–4 mm long, 4–8 mm wide; midrib keeled, extended as a straight, stout awn 5–8 mm long, apex acute; colleters in a line inside at the base of the stipule, 0.1–0.2 mm long, mixed with much longer simple hairs 0.2–0.5 mm long. Leaves on primary stem not seen; secondary stem leaves simple, opposite, equal, thickly leathery, matt, drying pale green above, whitish green below; petiole canaliculate 5– 10 mm long, 2–3 mm wide, densely hairy (hairs as stem), hairs to 0.5 mm long, Leaf blade elliptic, 8.8–10.5 × 4.7–6.3 cm, acumen 3 (– 7) mm long with rounded apex, base broadly acute to subtruncate, leaf edges a little decurrent on petiole, primary vein and secondary veins somewhat raised on the upper surface, clearly raised on the lower surface, with sparse brown hairs to 0.6 mm long, secondary veins 5–6 on each side of the midrib, arising at 40°– 50°, arching straight then towards margin, forming a weak, looping marginal nerve. Tertiary nerves inconspicuous and sparse. Tuft domatia between the midrib and secondary veins, domatia orbicular to longitudinally elliptic, 1 –1.5 mm long, with brown weakly crisped hairs 0.2–0.6 mm long. Inflorescence and flowers not seen. Infructescence axillary, 2.5–3.5 × 2– 3.5 cm, with 3–7 fruits, peduncle (2 –) 5 mm long, basal bract pair naviculate, 8 × 2 mm, apices awned; rachis bifurcate 1–4 mm from base, branches 5–13 mm long; bracts triangular, 2–3 mm long, 1–2 mm wide at base, apex acuminate to awned, inflorescence axes hairy, hairs brown, to 0.8 mm long. Fruits brown (from green) when ripe, dull, 1- or 2-seeded, bracteoles linear, c. 1 mm long, pedicel 2–3 mm glabrous; 1-seeded fruits ellipsoid 8–12 × 5–9 × 5–7 mm, calyx located at the side of the fruit, calyx 2–3 mm diameter, lobes 5–6, oblong-elliptic 0.4–0.6 × 0.1–0.3 mm, apex hooded, incurved (Fig 1 F) persistent; disc torus-like to saucer-shaped 0.6–0.8 mm diameter, drying glossy black, glabrous, surrounded by a ring of erect hairs, hairs 0.6 mm long; 2-seeded fruits heart-shaped (retuse and widest at apex, Fig 1 K), 10 × 11–12 × 6–7 mm, a little constricted between the carpels, calyx located in a sinus 1 mm deep between the carpels. Pyrene of 1-seeded fruit pale brown, ellipsoid or slightly reniform, lid shallowly convex, pointing sub vertically 1.5 × 2.5 mm crest shallow, with a longitudinal groove; pyrene of 2-seeded fruit ellipsoid, flattened along 1 side, 10.5 × 5.5 × 5.5 mm. Pyrene wall 0.15–0.3 mm thick, outer surface pale brown with low rounded projections separated by fibres; surface with glassy finger-print-like pattern (Fig. 1 K). Seed as pyrene, 7 × 5 × 4 mm, surface pustulate, dark brown, convoluted; seed in transverse section with endosperm tanniniferous areas dense, black, arranged in 12–14 rays (Fig 1.G), rays separated by bands of hard white endosperm, embryo cylindric, central.

#### Distribution

**GUINEA**. Forécariah Prefecture, southern plateau of Kounounkan Massif.

#### Habitat & ecology

Fissured sandstone rocks, among shrubs and small trees along a seasonal stream close to a sparsely wooded meadow at an altitude of 1100 m. The ecological conditions of this environment are influenced by the presence of seasonal water, which contributes to soil moisture and the diversity of surrounding plant species. In the submontane forest gallery where *Keetia kounounkan* was found, several associated plant species were observed,. These species include *Memecylon afzelii* G.Don, *Hibiscus kounounkan* Cheek ined., *Ternstroemia guineensis* Cheek, *Warneckea fascicularis* (Planch. ex Benth.) Jacq.-Fél., *Ficus ovata* Vahl, *Cailliella praerupticola* Jacq.-Fél., *Glenniea africana* (Radlk.) Leenh., *Kotschya uniflora* (A.Chev.) Hepper, *Keetia mannii* (Hiern) Bridson, and *Keetia susu*.

Individuals are typically scattered and associated with other flora adapted to similar conditions. The specialized nature of this habitat, however, makes the species vulnerable to environmental degradation, including fire and habitat fragmentation.

#### Conservation status

*Keetia kounounkan* is an endemic species from Guinea, currently known only from the southern plateau of the Kounounkan Massif in Forécariah Prefecture. Although it is represented by a single herbarium specimen, several other individuals of the species have been observed at the site.

Its extent of occurrence (EOO) is estimated to be no greater than 16 km^2^, based on the area of the southern plateau of the Kounounkan Massif from which the only known collection and observations have been made. Its area of occupancy (AOO) across the plateau area is also likely to be highly restricted but may narrowly exceed 10 km^2^. The plateau is inferred to represent a single location threatened by dry-season bushfires set by cattle herders. As a result of this threat, the species is inferred to be undergoing a continuing decline in habitat quality. The number of mature individuals cannot be reliably estimated, but it is suspected that the true value may exceed 1,000.

Given the availability of other similar submontane habitats in neighbouring Kindia and Dubréka Prefectures, it is possible that this species occurs at other sites; however, it has not yet been reported from collecting trips to neighbouring plateaux. Pending more precise data on its distribution and population size, and adopting a precautionary approach on the basis that its distribution may prove to be highly restricted, *Keetia kounounkan* is here provisionally assessed as Critically Endangered (CR) B1ab(iii), following IUCN criteria. Further survey work is essential to refine this conservation assessment; for example, confirmation of its presence on other plateaux may permit assessment at a lower category of extinction risk, though it would likely remain threatened.

Kounounkan was designated as a TIPA in 2019 (Couch et al. 2019) and is set to become a formally protected area. Of the 22 TIPAs in Guinea it has the highest number of strictly endemic species, with seven globally unique species recorded (Couch et al. 2019), including *Gladiolus mariae* Burgt (Iridaceae, van der Burgt et al. 2019), *Ternstroemia guineenis* Cheek (Ternstroemiaceae, Cheek et al. 2019). Subsequently, some of these species have been found elsewhere, but at the same time, additional new endemic and near endemic species have been published from Kounounkan and nearby sandstone plateaux e.g. the new genus *Benna alternifolia* Burgt & Ver.-Lib. (Melastomataceae, van der Burgt *et al*. 2022), *Ctenium bennae* Xanthos and *Trichanthecium tenerium* Xanthos (both Poaceae, Xanthos *et al*. 2020; 2021).

#### Etymology

The species is named after the Kounounkan plateau, as this is the only place where the species was observed. Kounounkan is of immense importance for plant conservation in view of the number of globally unique and highly threatened species present.

#### Notes

Among the species of *Keetia* found growing with *Keetia kounounkan, K. mannii* (Hiern) Bridson is morphologically similar but differs in having glabrous stems (or only a few hairs), leaves with an acute acumen apex, and pit domatia similar to those of *K. susu*. In contrast, *K. kounounkan* has densely pubescent young stems, leaves with a rounded acumen apex, and tuft domatia.

*Keetia kounounkan* is distinctive in the genus for its non climbing habit (resembling in this *Keetia susu*) and unusual also for the stiff bristle like hairs on the stems, petiole and abaxial veins of the leaf blade. The persistent stipules, with robust, long awns are also distinctive. The fruit colour, dull brown and matt when ripe is unusual in a genus where fruits are usually red or orange, sometimes black, when ripe. These features make the species readily identifiable despite its initial similarity to *K. susu*.

The apparent rarity of this species highlights the importance of continued research and conservation action to obtain it in flower, and to understand its full distribution, ecological preferences, and conservation needs. The clear morphological distinctions underscore the richness of the genus *Keetia* in Guinea.

The two specimens of the second species described in this paper, *Keetia tinka* (*Balde* 274 and *Fofana* 303) were formerly considered for inclusion as a subspecies of *K. magassoubiana* (formerly *K. sp. aff. tenuifolia* of Bridson, Cheek et al. 2025) until it was found that the number of morphological characters separating them, mainly qualitative (see Table 2) justified species rank. This conclusion is further supported by the different geographic and elevational ranges of the two taxa (Table 2). The two species also bear fruit (and so likely flower also) at different seasons. It seems highly probable that the two taxa are closely related due to the many similarities in all organs that are known for both species e.g. the fruits are almost indistinguishable.

**Table 2.** Diagnostic characters separating *Keetia magassoubiana* from *Keetia tinka sp. nov*. Data for the first species from Cheek *et al*. (2025) and specimens cited at K therein.

| Characteristic | *Keetia magassoubiana* | *Keetia tinka* |
| --- | --- | --- |
| Leaves of secondary stems: shape and base | Narrowly elliptic to oblong, length: breadth ratio 2.5–3.5: 1<br>Leaf base acute | Elliptic (rarely ovate-elliptic), length: breadth ratio 1.3–2(–2.25): 1<br>Leaf base obtuse to rounded |
| Domatia | Mainly present along midrib. Domed with a minute central aperture c. 0.1 mm diam., Hairs not visible within. | Mainly present at secondary nerve junctions. Open pits, aperture 0.25–0.5 mm wide. Hairs conspicuous within aperture. |
| Stipule awn | Stipule awn flat, straight | Stipule awn folded along midrib, arched |
| Peduncle bract pair | United at base, forming a sheath, margins lacinate | Free at base, not forming a sheath, each triangular, margins entire |
| Pyrene surface | Honeycombed (deeply pitted) | Subverrucate |
| Seed endosperm in transverse section | Conspicuous radial black tanniniferous bands | Tanniferous cells dispersed, inconspicuous, bands absent |
| Fruiting season | February to May | July to October |
| Elevational range (m) | 15–960 m | 1070–1380 m |
| Geographic range | Guinea (Guinée Maritime, Haute Guinée, Guinée Forestière), Ivory Coast, Sierra Leone, Liberia | Guinea (Moyenne Guinée) |

***Keetia tinka*** *Cheek & Simbiano* ***sp. nov***. Type: Republic of Guinea, Fouta Djalon, Dalaba Prefecture, Forêt Classée de Tinka, near Karéh, ‘Edge of secondary forest, disturbed area, old field’, 10° 22’ 50.0” N 12° 15’ 12.9” W, 1278 m, fr., 19 Oct. 2017, *Fofana F*. 303, with Larridon, I. Couch, C. & Haomou, A. (holotype K! barcode K000874709; isotype HNG). (Figure 4).

**Fig. 2.**
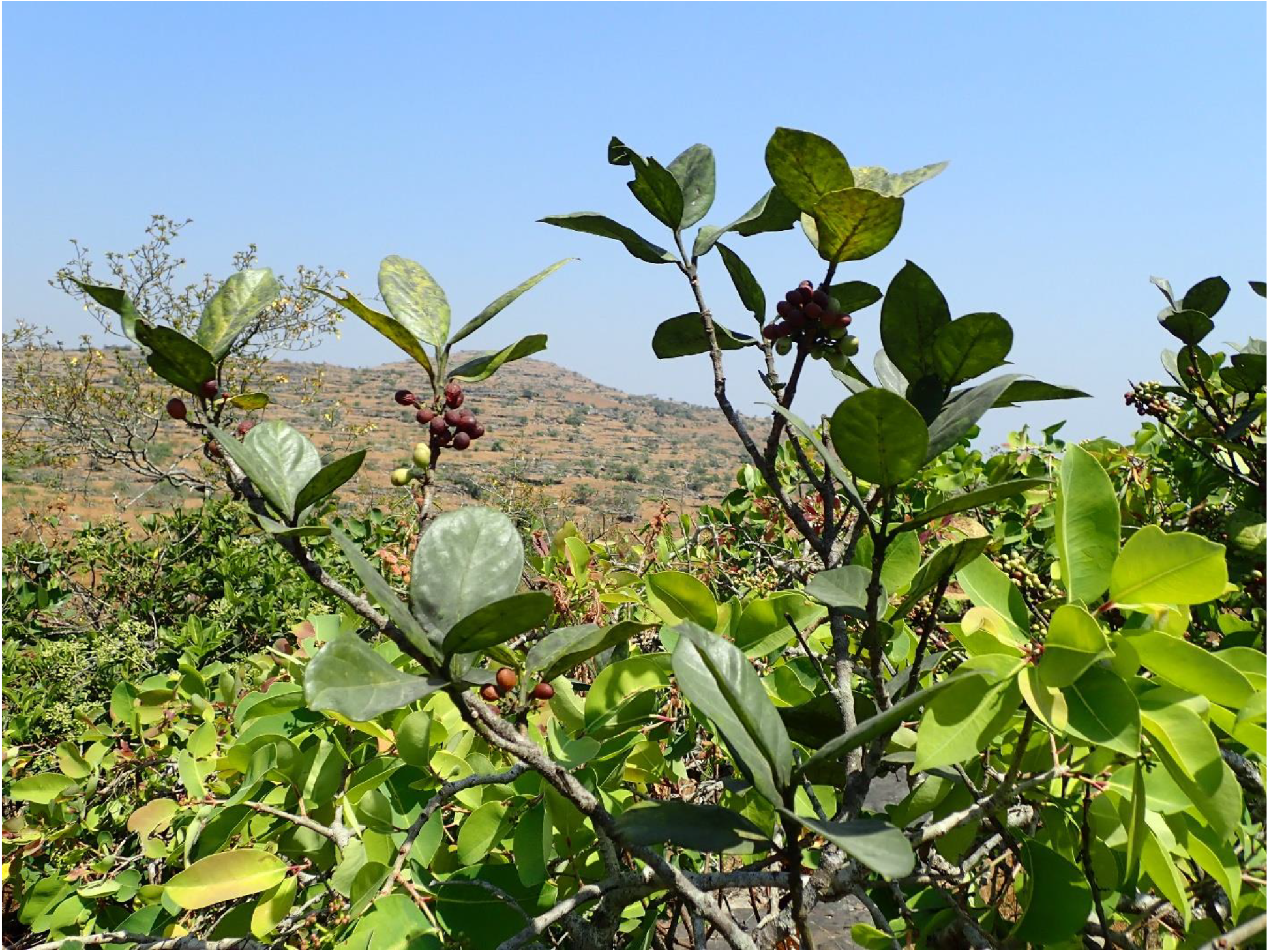
Keetia kounounkan. In habitat in stunted gallery forest of a seasonal stream set in grassland on the Kounounkan sandstone plateau. *Van der Burgt* 2262. Photo by Xander van der Burgt

**Fig. 3.**
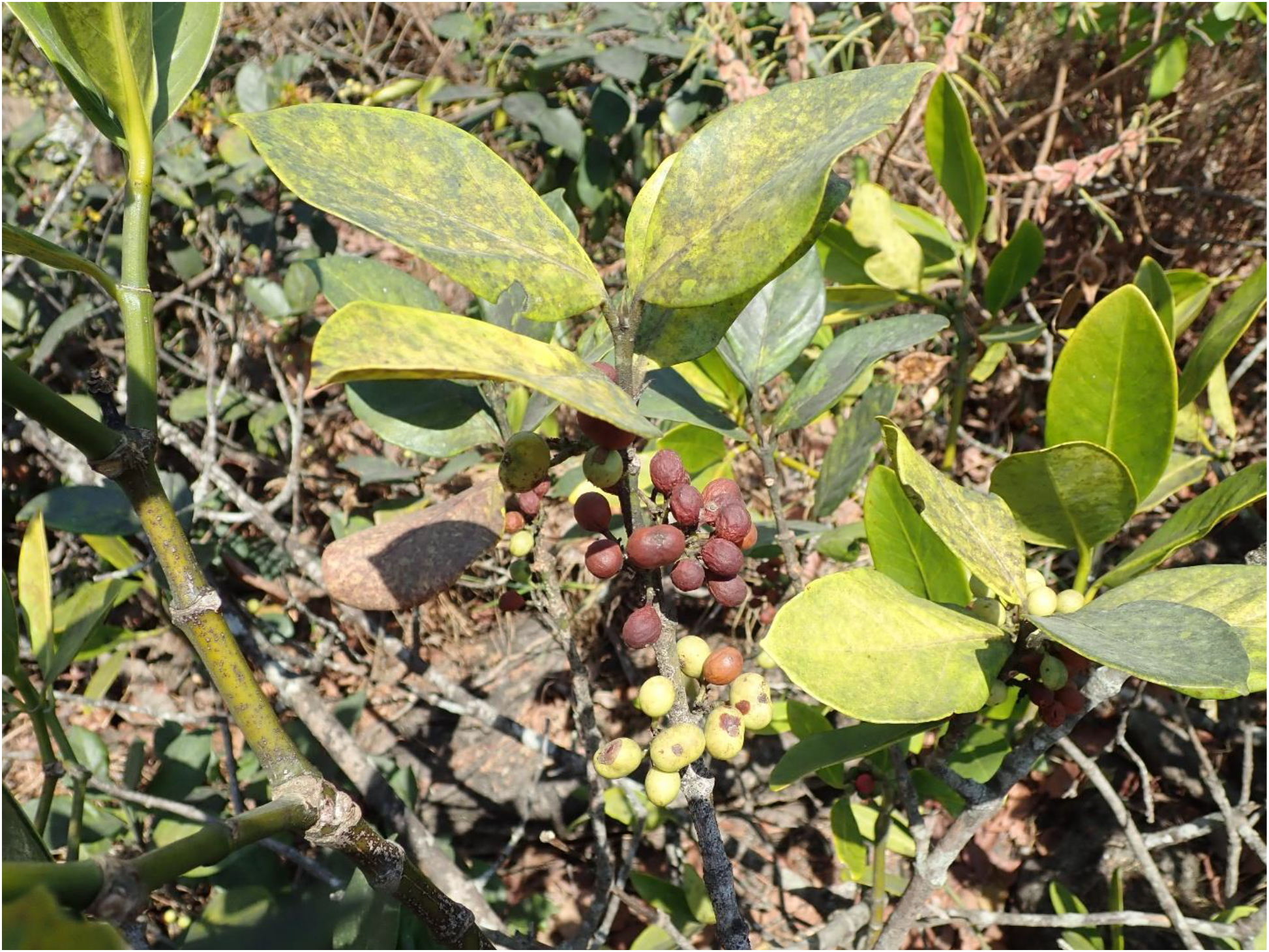
Keetia kounounkan. Close up showing the fruits, ripe and unripe. *Van der Burgt* 2262. Photo by Xander van der Burgt

**Fig. 4.**
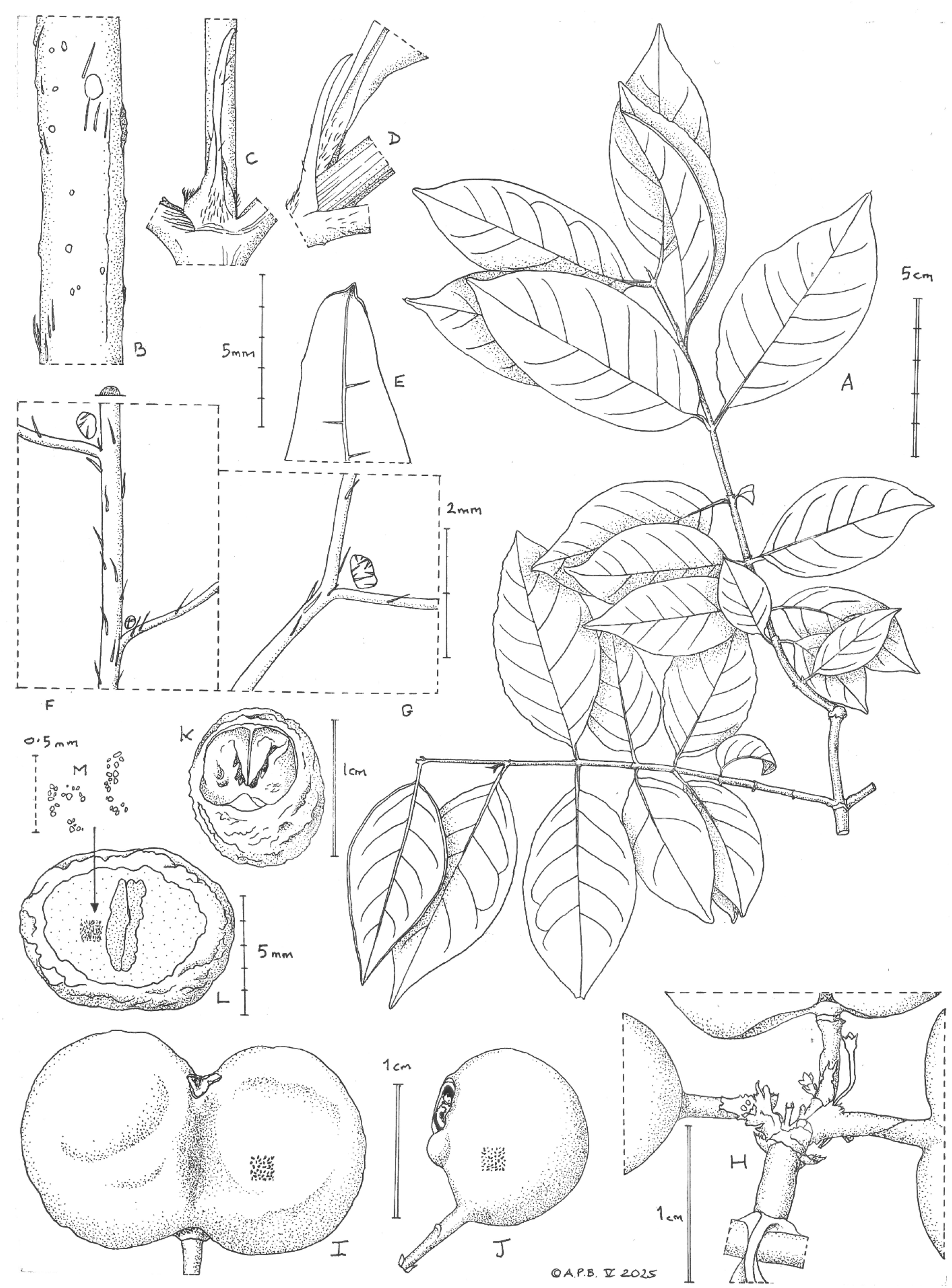
Keetia tinka. A habit, two leafy secondary stems; B secondary stem indumentum; C stipule, face view; D stipule side view, and petioles; E acumen with apical mucro; F midrib domatia (atypically present); G typical domatia of secondary nerve bifurcations; H infructescence axis; I 2-seeded fruit, side view; J 1-seeded fruit, side view; K pyrene showing the flat, ventral lid; L transverse section of seed with collateral cotyledons; M detail of the dispersed, inconspicuous tanniniferous cells of the endosperm. A-C, E-I, K from *Fofana* 303 (K), D, J, L-M from *Baldé* 274 (K). Drawn by Andrew Brown.

#### Diagnosis

*Keetia tinka* is similar to *Keetia magassoubiana* Cheek but differs in the leaf blades elliptic (rarely ovate-elliptic), length: breadth ratio 1.3–2(– 2.25): 1 and with leaf base obtuse to rounded (vs narrowly elliptic to oblong, length: breadth ratio 2.5–3.5: 1, leaf base acute). The domatia of *K. tinka* are mainly present at secondary nerve junctions, they are open pits, aperture 0.25–0.5 mm wide, with hairs conspicuous within the aperture (vs mainly present along midrib, domed, with a minute central aperture c. 0.1 mm diam. And hairs not visible within. The pyrene of *K. tinka* has a subverrucate surface and the endosperm in transverse section lacks tanniniferous rays (vs pyrene honeycombed, tanniniferous rays conspicuous).

#### Description

Lianescent evergreen forest shrub 5–6 m high. Primary stems cylindrical to slightly 4- angular, brown-black, internodes 1–3.5 × 0.2–0.5 cm, glabrous. Secondary stems (plagiotropic or brachyblasts), 12.5–17.5 cm long, with 6–9 nodes (Fig 4A), ascending, cylindrical, internodes 1.3–2.7 × 0.1–0.3 cm, hairs very sparse, slightly spreading, white, 0.4–0.6(– 1) mm long (Fig. 4B). Leaves of primary stems unknown, those of secondary stems simple, opposite, equal, blade thinly leathery, pale brownish-green to brownish-grey adaxially, abaxially pale whitish grey after drying, elliptic, less usually ovate-elliptic, (2.75–)3–8 × (1.3–)1.5–4 cm, apex acute to acuminate, with a short acumen 0.2–0.5(– 0.8) cm long, the acumen apex minutely mucronate and hooded (Fig 4E), base obtuse or rounded, sometimes slightly decurrent towards the top of petiole, asymmetrical, margins entire and slightly revolute. The adaxial (upper) surface with a raised midrib and secondary veins, abaxial (lower) surface bearing a few sparse, slightly spreading hairs. The secondary veins bright white, broad, 4 to 6 on each side of the midrib, arise at about 60°, bifurcating c. 5 mm from the margin, the branches uniting (abaxial surface) to form an inconspicuous, weak, looping inframarginal vein, tertiary veins rare and scarcely visible. Domatia absent at the junction of midrib and secondary veins, or rarely present at the most distal nodes (Fig 4F), frequent at the branches of secondary veins (Fig 4G), domatial pits, orbicular and c. 0.25 mm diam., or longitudinally elliptic, c. 0.5 × 0.25–0.3 mm, containing 5–12 straight orange hairs 0.1–0.3 mm long; indumentum of the midrib (both surfaces) and abaxial secondary veins moderately dense, with appressed, straight, stiff, acute, red-brown hairs, 0.3–0.5 (– 1) mm long; petiole canaliculate, (3–)4–7(– 8) × 0.8–1 mm with adpressed hairs 0.3–0.5 mm long.

Stipules more or less persistent until the 4–5 th node from stem apex, 2–7(– 8) × c.2 mm, base triangular, 2–2.2 × 2 mm, (including basal sheath c. 1 mm long) apical awn 5(– 6) × 0.3– 0.5 mm, folded in two along the midrib, arched (Fig 4D) apex acute or rounded; external surface with moderately dense hairs c. 0.25–0.3 mm long, adpressed, translucent; inner surface glabrous except for a line of colleters and hairs at the base. Colleters c. 5 per stipule, glossy brown or black, erect, conical 0.25 –0.3 × 0.1 –0.15 mm, exposed when stipules fall; hairs erect, wiry, red, 0.7–1 mm in length. Inflorescence and flowers not seen. Infructescences axillary on secondary stems at 2–3 successive or alternating nodes, 2.5–3.5 × 3.9–5 cm, 3–9-fruited, peduncle stout 3–12 mm long, bract pair opposite, inserted c. 2 mm below apex, triangular, 1.5(– 2) × 1.5(– 2) mm, moderately hairy, hairs grey 0.1– 0.3 mm long; rachis glabrous, bifurcate, the branches each bifurcating 2(–3) times, bracts and bracteoles slightly smaller than peduncular bracts. Fruits glossy dark brown after drying, surface smooth, glabrous; 2-seeded fruits (3 of 4 in *Fofana* 303, K) strongly didymous, with a deep furrow on both sides separating the globose carpels, 1.4–1.5 × 2.3–2.5 × 1–1.1 cm, apex and base emarginate, disc distinctly accrescent, 0.3–0.4 cm in diameter, V-shaped (due to carpel expansion), densely hairy, hairs translucent, straight, erect, 0.1–0.2 mm long. Calyx lobes erect, triangular, c. 0.6 × 0.5 mm, inner surface densely hairy, hairs c. 0.1 mm long, sinuous, thick. 1-seeded fruits (all 18 fruits in *Baldé* 274, K) as for 2-seeded fruits, but ellipsoid, 11–15 × 9–13 × 9–12 mm, pedicel attached obliquely, disc lateral, flat, often with an aborted carpel inserted between disc and pedicel; aborted carpel hemi-ellipsoid, c. 3 × 2 mm, 1.5 mm tall. Pyrene ellipsoid, 0.9–1.1 × 0.7–1 × 0.8–1 cm, apex and base broadly rounded, ventral surface slightly convex; lid ventral, nearly flat, semicircular, 6.5 × 7.5 mm, crest indistinct with a cleft along the midline; wall, c.0.5 mm thick; outer surface subverrucate, inner surface smooth and shiny. Seed ellipsoid 9 × 5–5.5 × 4–6 mm, surface convoluted, brain-like, black-brown, in transverse section endosperm with thinly dispersed and inconspicuous tanniniferous cells (bands absent); embryo with two flat cotyledons (Fig. 4K).

#### Distribution

Guinea, Fouta Djalon, Dalaba Prefecture, Forêt classée de Tinka and Tangama.

#### Additional specimen examined

**REPUBLIC OF GUINEA**. Fouta Djalon, Dalaba Prefecture, Commune Urbaine Dalaba, pres de Yomou. Foret classee de Tangama, 10° 40’ 25.8” N 12° 15’ 49.6” W, 1328 m elev., fr., 12 July. 2017, *Baldé, A*. 274 with Couch, C., Hooper, F., Kouliye, M. & Diallo, M. (HNG, K).

#### Habitat & ecology

Submontane forest and edge of secondary forest. Elevation: 1070–1380 m (elevations read from Google Earth).

#### Conservation status

*Keetia tinka* is a species on current evidence endemic to the Fouta Djallon of Guinea, where it is known only from two sites, the Tinka Classified Forest, near Dalaba, and c. 30 km to the South, west of Mamou, in the Tangama Forest. Between these two sites the original submontane forest habitat is extremely fragmented to non-existent as it is throughout the Fouta Djalon, due to extensive and intensive clearance for agriculture over recent centuries: “Submontane forest with threatened species has been all but eliminated from the `core’ Fouta Djallon area that extends from Mamou, north to Dalaba….Those forest reserves that survive, such as the Tinka Classified Forest near Dalaba, have been heavily managed for forest products and appear to have lost the higher-level threatened species that they probably once possessed. Efforts to rediscover such species…..have so far failed.” (Couch et al. 2019 p. 54). The Tinka forest site has been heavily managed for production (and not nature conservation) as stated above, even though the canopy is intact, and these threats continue (Cheek & Couch pers. Obs. 2016 onwards, during extensive surveys of submontane forest in the Fouta Djalon with HNG teams). The Tangama forest is much more heavily disturbed than Tinka, with fields of cultivation (noted on the specimen label), and even in Google Earth Pro imagery dated 2024, about half of the area around the collecting point for our specimen is still cleared and lacks trees entirely. Threats to the habitat from agriculture continue. Therefore, two threat-based locations can be accepted

The area of occupancy (AOO) is 8 km^2^ using the stipulated IUCN grid cells. The extent of occurrence (EOO) cannot be calculated from two points, but we take to be as the same as the AOO. *Keetia tinka* can therefore be provisionally assessed as Endangered (EN) B1ab(iii)+2ab(iii) according to IUCN 2012 criteria. Further studies are needed to better understand the ecology, population and threats and threats for this species. It is to be hoped that further sites might be found for the species. In the meantime species conservation posters for the species should be made and deployed to sensitise local communities in the vicinity of this species as to its importance. Efforts should also be made to collect seed for possible conservation, but also for immediate propagation to attempt to multiply the species at safe sites to reduce the risk of global extinction.

#### Etymology

The species is named after the Tinka forest, Dalaba, as this is where the species appears to have the best possibility of surviving.

#### Phenology

Fruiting July–Oct.

#### Notes

The two specimens of this species, *Balde* 274 and *Fofana* 303 were formerly considered for inclusion as a subspecies of *K. magassoubiana* (formerly *K. sp. aff. tenuifolia* of Bridson) until it was found that the number of mainly qualitative morphological character separating them (see Table 2) justified species rank. This conclusion is further supported by the different geographic and elevational ranges of the two taxa (Table 2). The two species also bear fruit (and so likely flower also) at different seasons.

## DISCUSSION

The publication of these two new species of *Keetia* will increase the total for Guinea from the 10 *Keetia* species previously recorded (Gosline et al. 2023b), of which two are endemic, to 12 and four species respectively. This exceeds the total published for Gabon (Sosef et al. 2006) despite this being considered a far more species-diverse country. These numbers help to illustrate the unexpectedly high species diversity of Guinea and the progress now being made towards completing its inventory.

The main, most contiguous and most well-known part of the sandstone plateaus of Guinea are the Fouta Djalon that dominate Moyenne Guinee. This area is densely populated and natural habitats have been heavily impacted. Many threatened plant species recorded there a century ago have not been refound despite targeted searches (Couch et al. 2019). The discovery of a new taxon to science there (*Keetia tinka*), from recent collections, is therefore unexpected and gives hope that with further surveys, more threatened taxa might be found than are known now, even in non-pristine, secondary areas.

In contrast to the Fouta Djalon proper, the sandstone plateau to the west, closer to the Atlantic, are less densely inhanbited and probably for this reason continue to provide a flow of new species and even genera to science, despite being negatively affected primarily by grazing and artificial fires. While the southern part of this block that includes Kounounkan has seen the largest part of these discoveries, the northern part, around Kindia has also yielded discoveries, e.g. the new genus *Kindia* Cheek (Cheek et al. 2018b), and species such as *Tephrosia kindiana* Haba, B.J.Holt & Burgt (Leguminosae, Haba *et al*. 2023). These are summarized in the paper describing *Virectaria stellata* Cheek et al. (Rubiaceae, Simbiano et al. 2024).

About 75% of plant species new to science published today are already threatened (Brown *et al*. 2023). Usually this is because they have small ranges making them at risk of extinction from habitat clearance, making description urgent so that they can be Red Listed if this is merited, and prioritized for conservation action (Cheek *et al*. 2020b). Conservation actions such as improved selection and prioritization of areas for conservation (Darbyshire et al. 2017) are needed if species such as those described in this paper are not to become globally extinct as have so many other plant species (Humphreys *et al*. 2019;) This is especially urgent in Guinea where over 90% of original forest habitat was considered lost before the end of the 20^th^ century (Sayer et al. 1993) and that which survives is fast being cleared. Fortunately there are positive indications that most of the area of Guinea prioritized as Important Plant Areas by Couch et al. (2019) will receive support for biodiversity protection in the near future.

## ACKNOWLEDGEMENTS

The authors thank the funders of the Guinea TIPAs programme for enabling this paper to be developed, especially JRS Biodiversity for supporting the first author training on this taxonomic research project, but also the Darwin initiative capacity building fund, the Dinswade Trust for our field and plant species conservation work with local communities in Guinea and the Franklinia Foundation for supporting “Conservation of threatened trees species in three Tropical Important Plants Areas of Guinea”, and the Darwin Initiative of the Department of the Environment Food and Rural Affairs (DEFRA), UK government (project Ref. 23–002). Mr Abdoulaye Yéro Baldé, former Minister, Guinean Ministry of Higher Education and Scientific Research, Dr Binko Mamady Touré, former Secretary General of the same Ministry, and Dr. Facinet Conté, Secretary General of the same Ministry, are thanked for their cooperation. Colonel Layaly Camara, former Director, Direction National des Eaux et Forêts, Mr Mamadou Bella Diallo, Nana Koulibaly, T. Delphine Kolié, and Mr Alpha Illias Diallo, CITES Focal Point, Direction National des Eaux et Forêts, authorised the export of the plant specimens. The first author’s training visit to Kew to write this paper was funded by the JRS Biodiversity grant (70022) “Enhancing data access to transform Guinea’s capacity to identify and protect its threatened plants”. The Prefects of Forécariah and Kindia Prefectures are thanked for their hospitality during the fieldwork. Two anonymous reviewers are thanked for constructive comments on an earlier draft of the paper.

